# Benchmarking Long-read Sequencing Tools for Chromosome End-specific Telomere Analysis

**DOI:** 10.1101/2025.01.29.635538

**Authors:** Jake Reed, Mark Oelkuct, Kevin Coombes

## Abstract

Measuring chromosome end-specific telomeres is of great importance and could help elucidate better treatment algorithms and aid in a better understanding of cancer, aging, cardiovascular disease, and neurodegenerative diseases. In this study, we present a comparison of two cutting edge long-read sequencing telomere length analysis tools, TECAT and Telogator. We perform a comprehensive bench-marking of these two tools using Telseq as the standard. Our analysis included evaluating these tools on sensitivity, accuracy, and computational efficiency using a diverse data set of 9 samples from the 1000 Genomes Project which have matched long-read and short-read sequencing data. We found that while Telogator demonstrated superior sensitivity, identifying on average 31% more telomeric reads across all samples, TECAT showed better accuracy with measurements more closely aligned with established literature values and Telseq benchmarks (R² = 0.74 vs 0.37), and TECAT displayed better computational efficiency, completing tasks approximately 41% faster. Both tools successfully mapped telomere lengths to individual chromosome arms, demonstrating unprecedented resolution for telomere length analysis. Our results provide crucial insights for researchers selecting tools for telomere length analysis and highlight the current capabilities and limitations of computational approaches in telomere biology.

## 1 Introduction

Telomeres, the specialized nucleoprotein structures at chromosome ends, play crucial roles in maintaining genomic stability and cellular aging [1, 2]. The length of telomeres serves as a critical biomarker for cellular senescence, aging-related diseases, and cancer progression [3, 4]. While conventional methods like Terminal Restriction Fragmentation (TRF) analysis and quantitative PCR provide average telomere lengths across all chromosomes [5, 6], these approaches mask the biological significance of individual chromosome end telomere lengths, which can vary significantly and may have distinct functional implications [7–9]. For example the range of lengths found by TECAT for HG002, range from 3.39 kb to 7.19 kb and for Telogator from -0.90 kb to 10.69 kb. These methods all have their strengths and weaknesses, but as of yet, there are only two developed tools for measuring chromosome end-specific telomeres, TECAT ant Telogator.

Recent technological advances have led to the development of several methodologies capable of measuring telomere lengths at specific chromosome ends. These include Single Telomere Length Analysis (STELA) [10], Telomere Shortest Length Assay (TeSLA) [11], and Universal STELA (U-STELA) [12], each with its own strengths and limitations. Additionally, newer approaches use next-generation sequencing [13] and specialized bioinformatics tools [14] have emerged, these tools measure average telomere length across an entire sample use NGS. Third-generation sequencing offers the potential to directly sequence telomeres, providing single-molecule resolution of telomere lengths [7–9, 15, 16].

Several tools have become available which have the ability to computationally measure telomere lengths at the single-molecule resolution. Telomere length determination (TLD) was the first long-read sequencing tool to accomplish this in *Plasmodium falciparum* [15] followed by Telogator [16] in *Homosapiens*. TECAT, the successor to TLD and the tool being used for analysis in this study, has several updates compared to TLD, including being capable of analyzing human samples and a graphical user interface (shiny app) for generating figures from the results. These tools have the ability to measure telomere lengths at the single-molecule resolution. This opens a new avenue for researchers to study telomere lengths at the single-molecule resolution which will allow for the generation of completely novel hypotheses.

Despite the availability of these diverse methodologies, a systematic comparison of their accuracy, sensitivity, technical requirements, and practical applications remains lacking [17, 18]. Such comparative analysis is essential for researchers to make informed decisions about which method best suits their specific research questions and laboratory capabilities. Furthermore, understanding the complementarity between different approaches could enable more robust experimental designs that leverage the strengths of multiple methods [19].

This study presents a comprehensive comparative analysis of current tools and methodologies for measuring chromosome end-specific telomere lengths. We evaluate these methods across multiple parameters, including but not limited to: detection range, sensitivity to short telomeres, technical complexity, cost-effectiveness, and throughput capabilities [12, 20]. Our analysis aims to provide a framework for method selection while highlighting areas where further technological development may be needed to address current limitations in telomere length measurement. All of these analyses are based on mean telomere length instead of chromosome end-specific telomere lengths. This is due to the limitation of the current academic literature lacking in the chromosome end-specific lengths. This is mainly due to the fact that these tools are at the cutting edge of technology and have not been widely adopted yet, and there are no other known methods for measuring chromosome end-specific telomere lengths.

## 2 Results

We collected data from the NCBI sequence read archive (SRA) for 9 samples from the 1000 Genomes Project. The samples were chosen because they included both whole-genome long-read and short-read sequencing data. The samples were processed and analyzed using a variety of computational tools, including BWA [21], samtools [22], minimap2 [23], TECAT, Telogator [16], Telseq [24], snakemake [25], R [26], plyr [27], knitr [28], ggplot2 [29], and ggpubr [30] to name a few. All sample IDs and accession codes are available in Table 1. We ran the tools on these samples and analyzed them for sensitivity, accuracy, throughput, and cost. The results of the analysis are presented below.

**Table 1.**
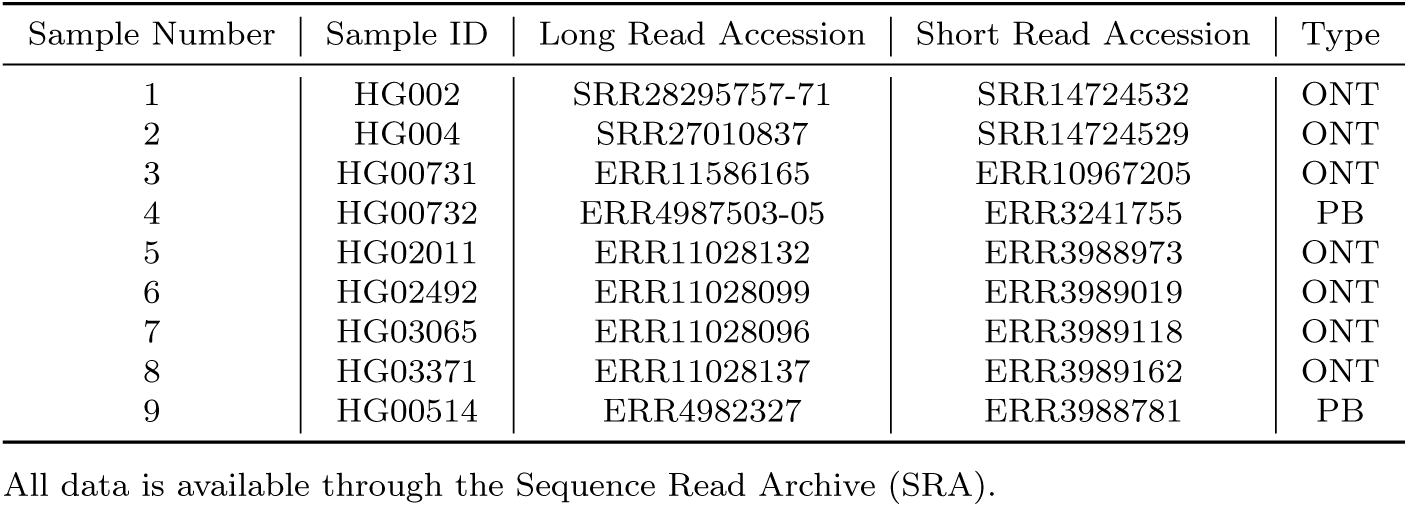
Summary of samples.

### 2.1 Sensitivity and Accuracy of Mean Telomere Length

#### 2.1.1 Sensitivity

The accuracy of computational tools for chromosome end-specific telomere analysis is heavily influenced by the limitations of current sequencing technologies, both oxford nanopore and PacBio. Tan et al. highlight the prevalence of extensive basecalling errors at telomeric repeats, which hinders precise interpretation of telomere sequences across various organisms [31]. Such errors can lead to mischaracterization of telomere structure, negatively affecting downstream analyses and conclusions about telomere function in processes like aging and cancer. While PacBio has better per base accuracy, the reads are often shorter, and need special DNA extraction and shearing protocols in order to get the most accurate results [15]. Moreover, the inherent complexity of telomere variations, such as neotelomeres and chromosomal fusions, underscores the need for robust computational frameworks to accurately analyze these alterations [32].

Telomeric content in whole-genome sequencing data can be estimated with a simple calculation (e.g., number of ends times average telomere length divided by total human genome size). This results in a telomeric content of human whole-genome sequencing data to be approximately 0.017% (46 ends * 11.7 kb [33] / 3.27e6 kb [34]). This is a very crude estimate, but it does demonstrate the relative sparseness of telomeric repeats in whole-genome sequencing data. This presents unique challenges for computational tools due to the low signal-to-noise ratio in such datasets. As telomeres play a pivotal role in maintaining genomic stability, the development of more sensitive computational approaches is essential to improve telomere length measurement accuracy and better elucidate their implications in health and disease [35]. Advancing these tools is critical for enhancing our understanding of telomere biology and its relevance in clinical research and applications. For HG002, we find that Telogator identified 0.048% of the reads as telomeric, while TECAT identified 0.036% of the reads as telomeric which above the estimated value (0.017%) of the human genome being telomeric. This suggests that both tools are realiably identifying telomeric reads in the data while Telogator seems to excel at identifying telomere reads.

Telogator outper-formed TECAT in sensitivity across all samples, with a mean of 31% more reads identified as telomeric. The difference in sen-sitivity between the two tools was most pronounced in samples HG002 and HG004, where Telogator iden-tified 34% and 76% more telomeric reads, respectively. The dif-ference in sensitivity between the two tools was least pronounced in sample HG00732, where Telogator identified 3% more telomeric reads. These results suggest that Telogator is more sensitive than TECAT in identifying telomeric reads in long-read sequencing data (Fig. 1).

**Fig. 1.**
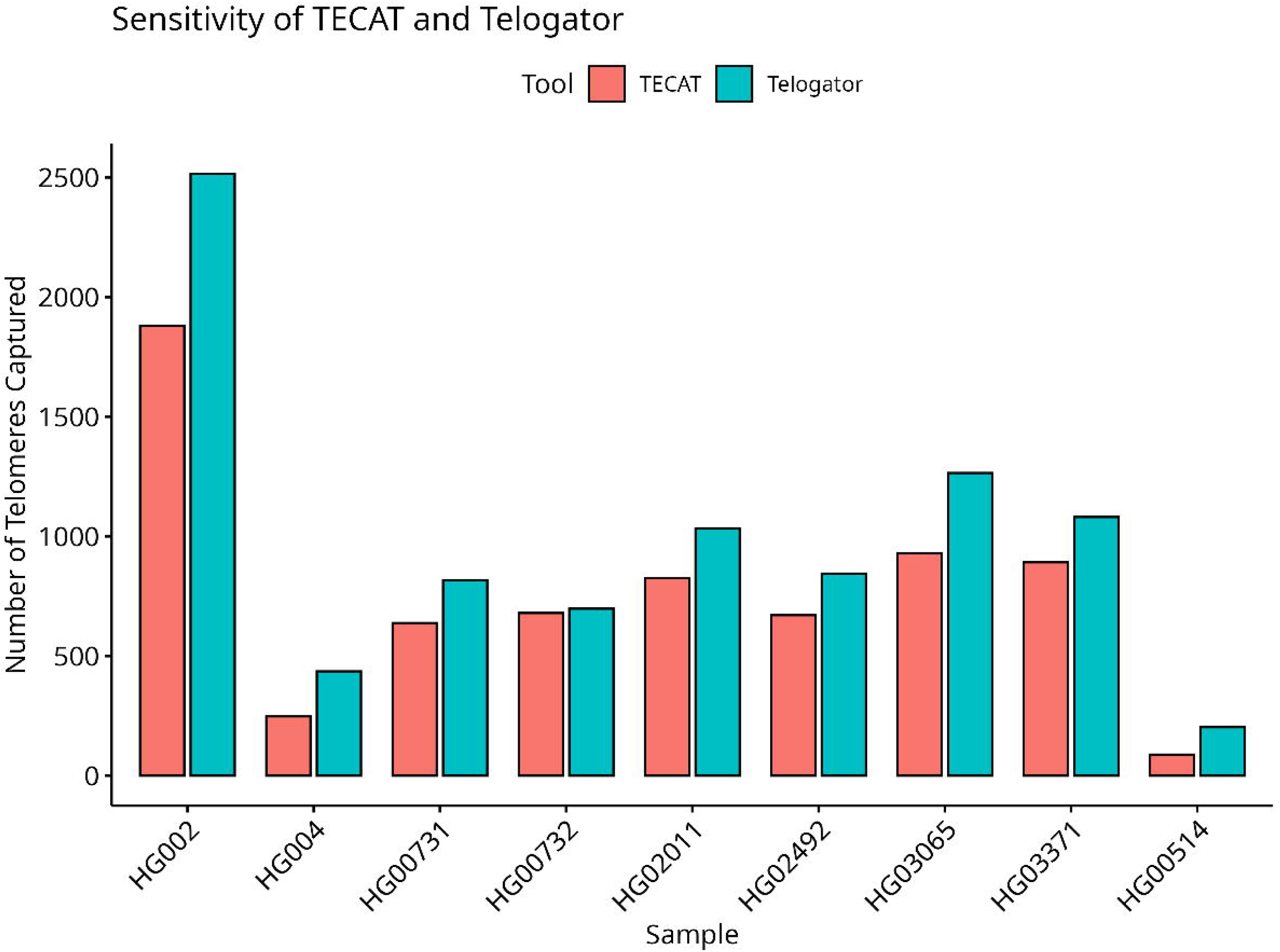
Raw number of telomere reads which were isolated from sequences.

#### 2.1.2 Accuracy

For accuracy, both TECAT’s and Telogator’s calculated mean telomere lengths fell within the general range of values reported in the literature for terminal length fragmentation experiments, which is approximately 4 to 6 kilobases (kb) [9] (Fig. 2). TECAT’s mean telomere length for sample HG002 was determined to be 5067 base pairs (bp) with a standard deviation of 1937 bp, closely aligning with the literature-reported telomere length of 5247 to 5270 bp [9]. Telogator, in comparison, calculated a mean telomere length for HG002 of 3503 bp with a standard deviation of 2050 bp, which was lower than the literature-reported range. The difference in mean telomere length between the two tools was most pronounced in sample HG002, where TECAT’s mean value was within 200 bp of the literature value, while Telogator’s estimate was approximately 2000 bp shorter (Fig. 2). These results suggest that while both tools provide estimates within the expected range for terminal length fragmentation experiments, TECAT produced values closer to the literature-reported mean for sample HG002. Both tools demonstrate the utility of computational approaches for telomere analysis, with TECAT showing an advantage in this particular dataset by producing estimates more consistent with the expected values.

**Fig. 2.**
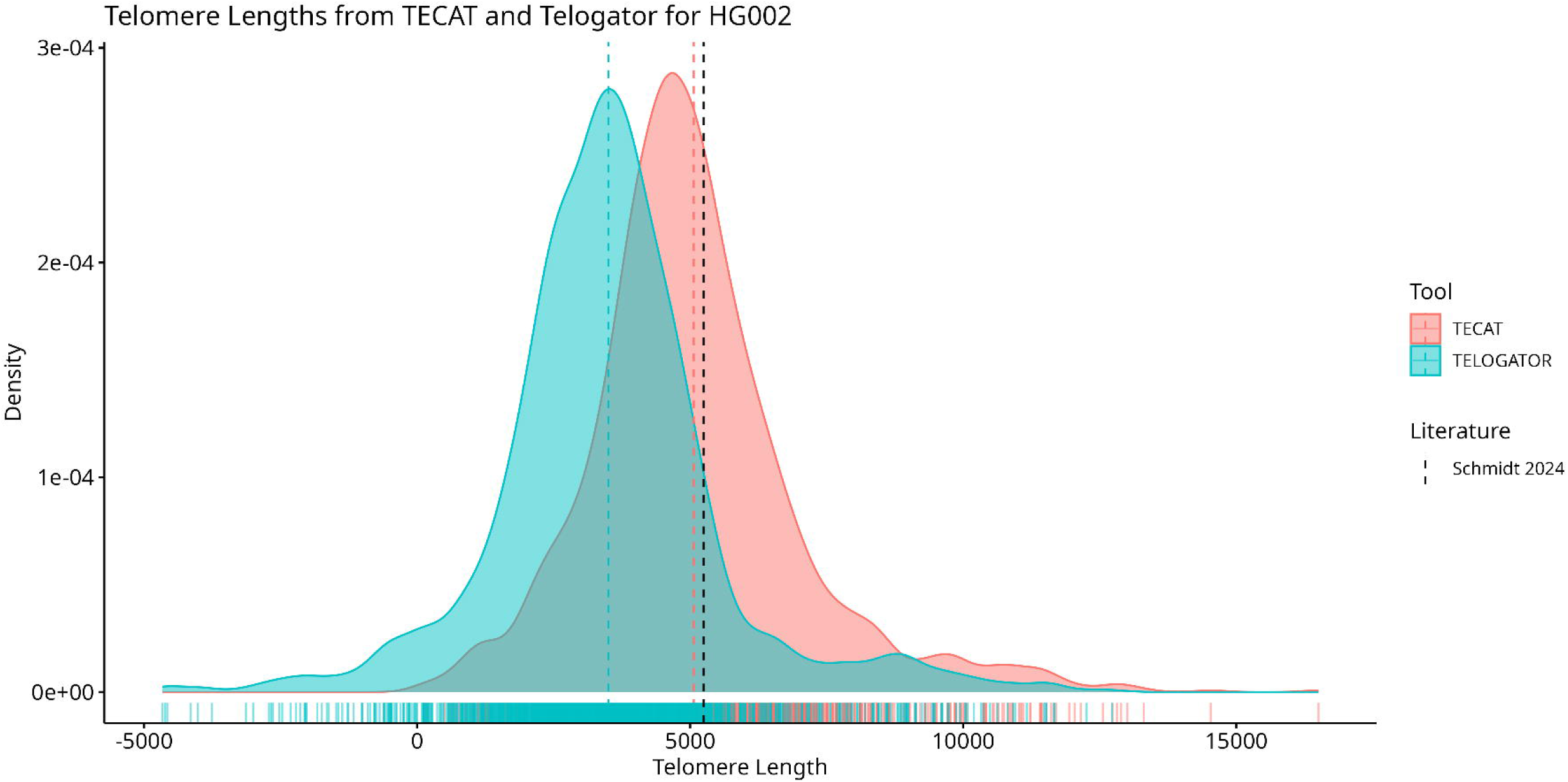
Density histogram of the telomere lengths from TECAT and Telogator as compared to literature values for telomere length in HG002

It also appears that Telogator is reporting negative telomere length values. I believe if this were to be fixed, the accuracy of Telogator may improve drastically. Every attempt was made in order to ensure that the data was processed correctly, and the correct parameters were used for each tool. There is no parameter that could be changed which would fix the negative telomere length values. This is a limitation of the tool and should be addressed in future versions.

Telseq is a tool that uses sequencing composition of telomere repeats in sequencing data in order to estimate average telomere length across the whole sample. The aggregate difference in telomere length between TECAT and Telseq was 16.49 kb while the aggregate difference between Telogator and Telseq was 34.98 kb (Fig. 3). The mean difference in telomere length between TECAT and Telseq was 1.83 *±* 1.90 kb while the mean difference between Telogator and Telseq was 3.89 *±* 2.27 kb (Fig. 3B). In every sample tested, TECAT was closer to the Telseq value than Telogator was to the Telseq value (Fig. 3A). This suggests that TECAT is more accurate in measuring telomere length than Telogator.

**Fig. 3.**
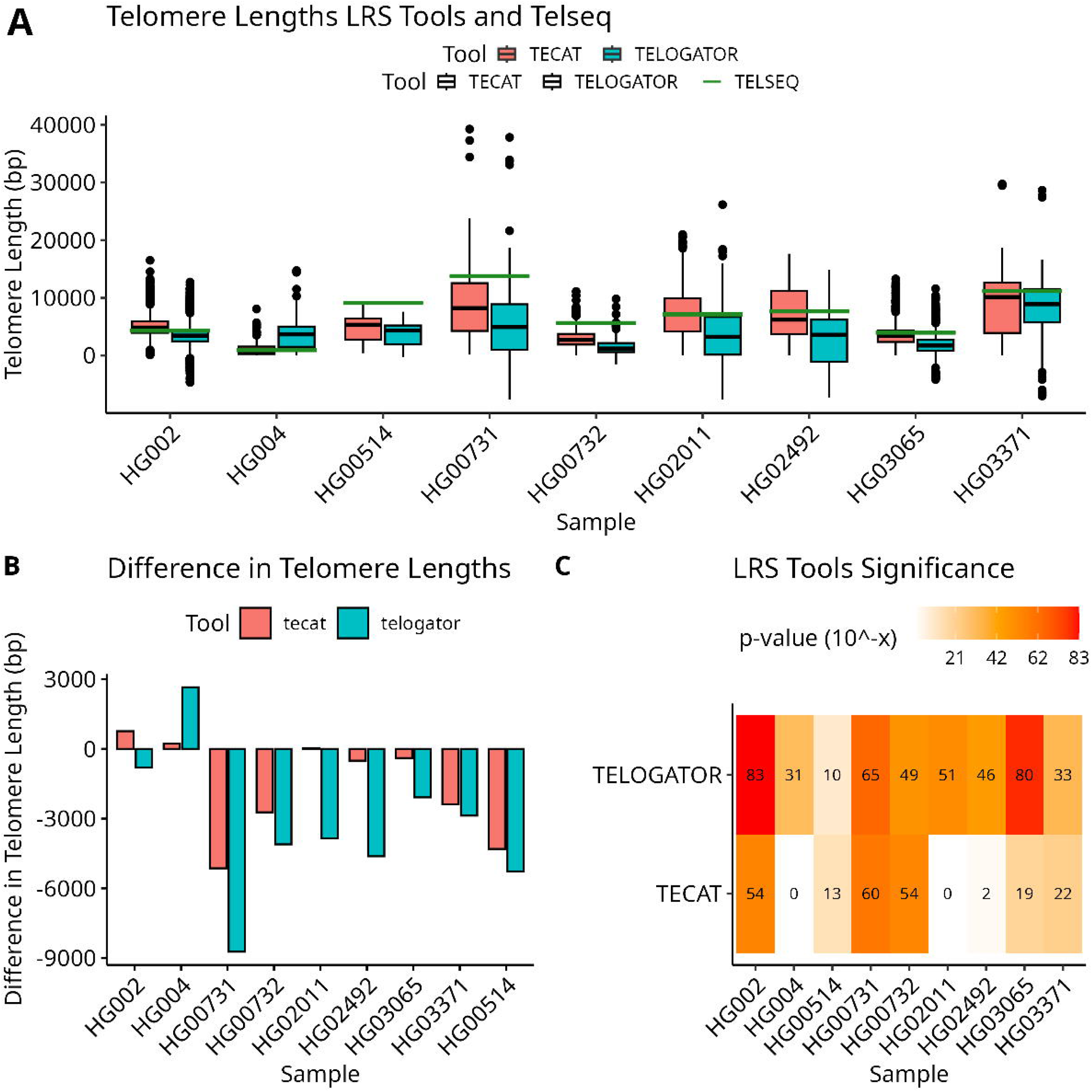
Differences in telomere length across all samples. A) Shows the length distributions as box-plots with red, blue and green representing TECAT, Telogator and Telseq, respectively. B) Shows the difference between TECAT and Telogator with regards to the Telseq measurement. C) heatmap where the colors correspond to statistical significance of a Wilcoxon test with white being non-significant and purple being highly significant. The values are the negative base-10 logarithm of the p-value.

For a statistical analysis one-sample Wilcoxon tests were performed on the differences between TECAT and Telseq and Telogator and Telseq. The results of the tests are shown in Fig. 3C. TECAT had fewer significant differences than Telogator with 3 samples showing no significant differences while Telogator did not have a single sample which was non-significant as compared to Telseq. In Figure 4B, a two-sample wilcoxon test was performed on the aggregate telomere lengths of each tool and the Telseq data. TECAT’s distribution of telomere lengths was not significantly different from Telseq’s distribution (p-value = 0.24) while Telogator’s distribution was significantly different from Telseq’s distribution (p-value = 0.011). This suggests that TECAT is more accurate in measuring telomere length than Telogator.

**Fig. 4.**
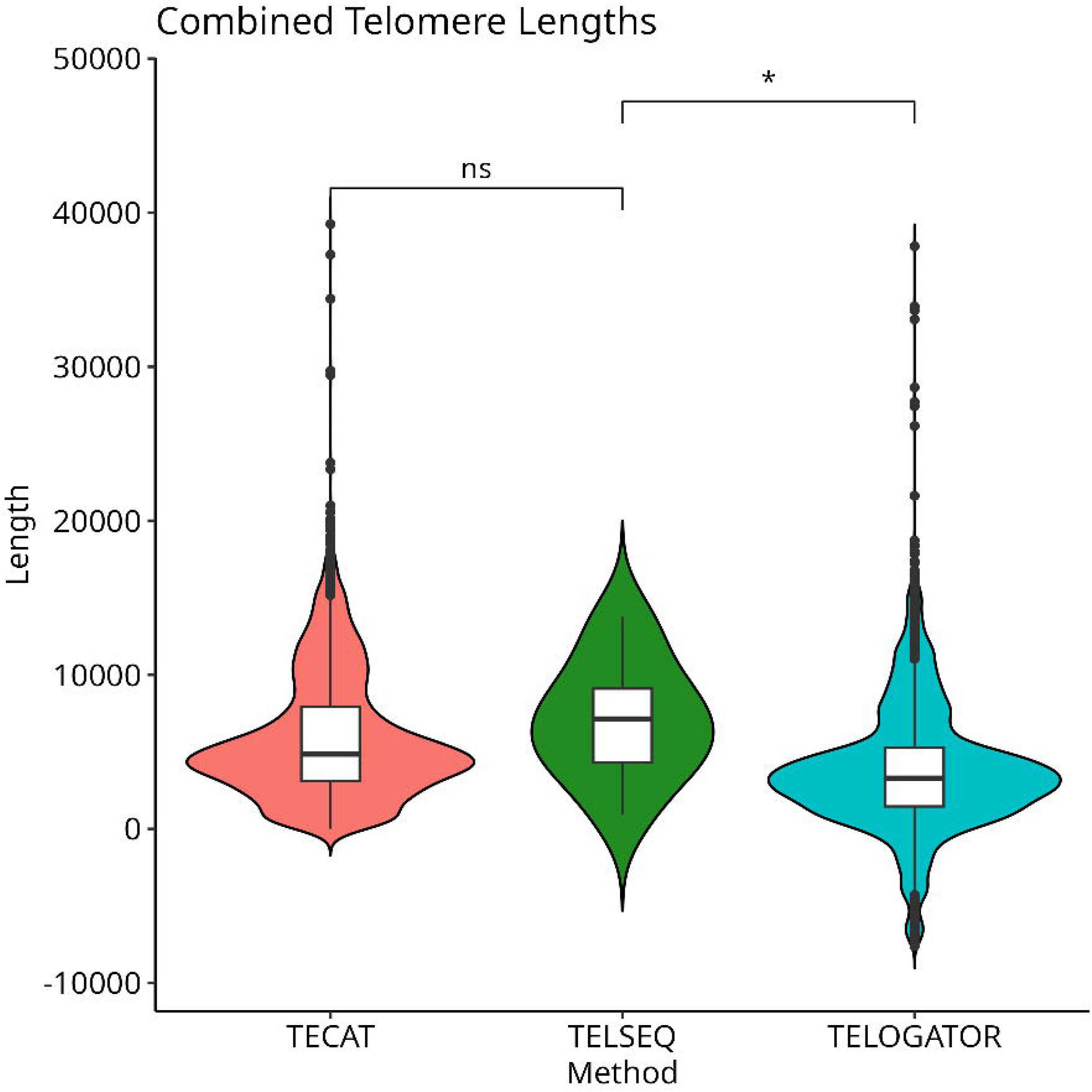
Statistical differences in telomere length. Shows the aggregate telomere lengths based on tool with the asterisk representing a Wilcoxon test results.

Linear regression was performed in R using least squares regression on the mean telomere lengths of TECAT and Telogator compared to the mean telomere lengths of Telseq. The R-squared value for TECAT was 0.74 while the R-squared value to Telogator was 0.37 (Fig. 5). Overall, TECAT was more accurate than Telogator. This suggests that TECAT is more accurate in measuring telomere length than Telogator.

**Fig. 5.**
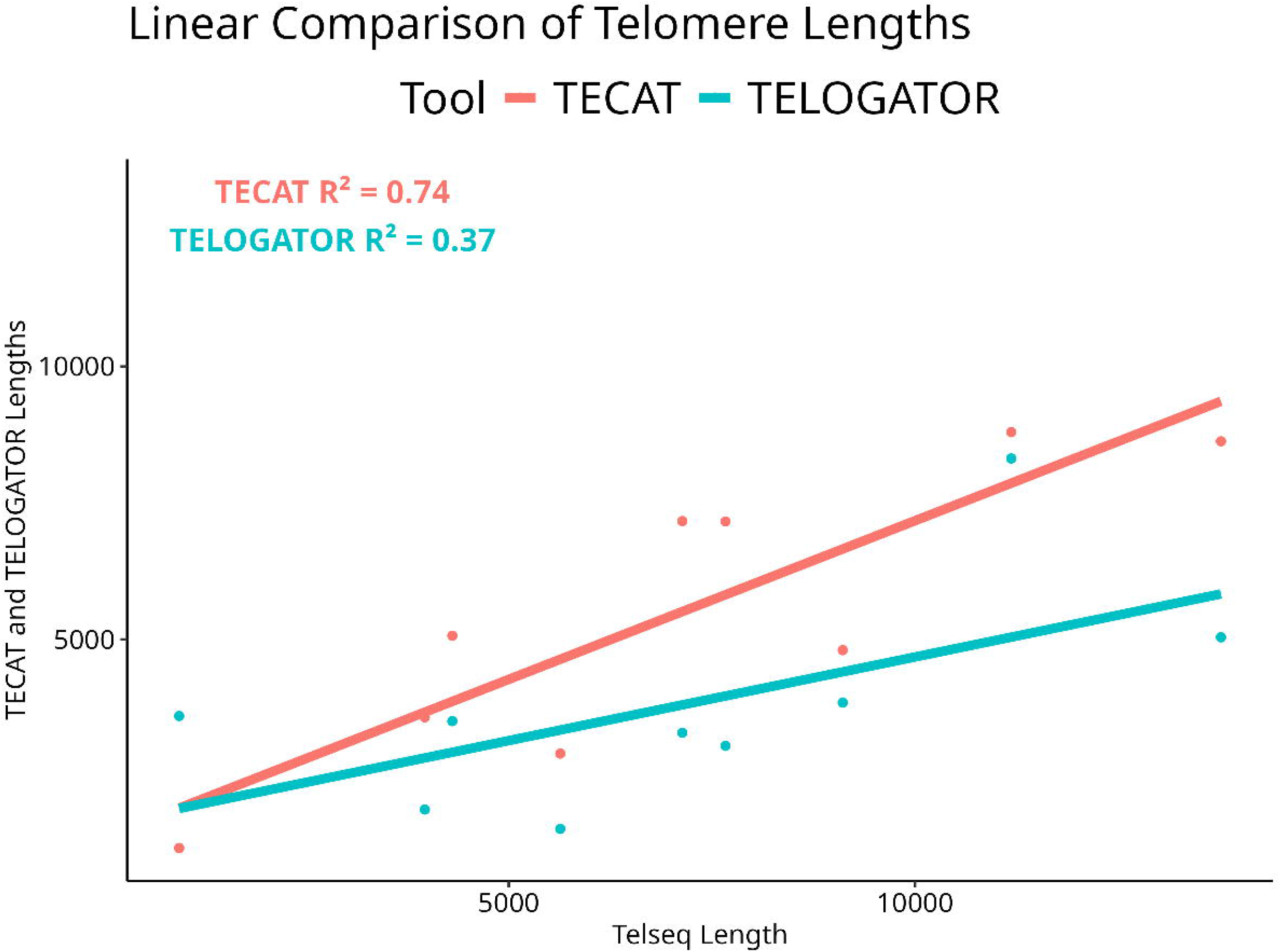
A linear comparison of telomere lengths based on tool. The x-axis is the mean telseq length for each sample and the y-axis is the mean TECAT or Telogator length which is annotated with a red and blue line, respectively. Linear regression was performed and the R-squared value is noted on the plot.

### 2.2 Throughput and Cost

For each sample processed, 18 threads of an i7-12700H CPU at 2.2GHz were used in order to process each of the samples, and the total run time for each of the tools was recorded. The total run time for TECAT for all samples was 829 minutes while Telogator had a total run time of 1398 minutes. The mean run time per sample of TECAT was 92 *±* 61 minutes and 155 *±* 114 minutes for Telogator. While this is a very useful metric a better gauge of efficiency and performance is megabytes processed per second which gives a more thorough presentation of efficiency.

TECAT outper-forms Telogator again, but both tools per-form adequately and are able to process most samples within 3 hours using 18 threads. When com-paring the mean time taken to process each sample, TECAT is 41% faster than Tel-ogator; however, when you normalize the data to megabases processed per second, the margin shrinks to 8% (Fig. 6). Figure 6 shows that TECAT processed a median of 14.63 ± 20.03 MB/s and Telogator processed a median of 10.36 ± 23.62 MB/s (median ± IQR). This suggests that TECAT is slightly more computationally efficient than Telogator. The i7-12700H consumes 45 watts per hour. An estimate of cost based on kilowatt hours would be $0.08/sample and $0.14/sample or $0.13/100GB and $0.21/100GB for TECAT and Telogator, respectively. This suggests that TECAT is more cost-effective than Telogator.

**Fig. 6.**
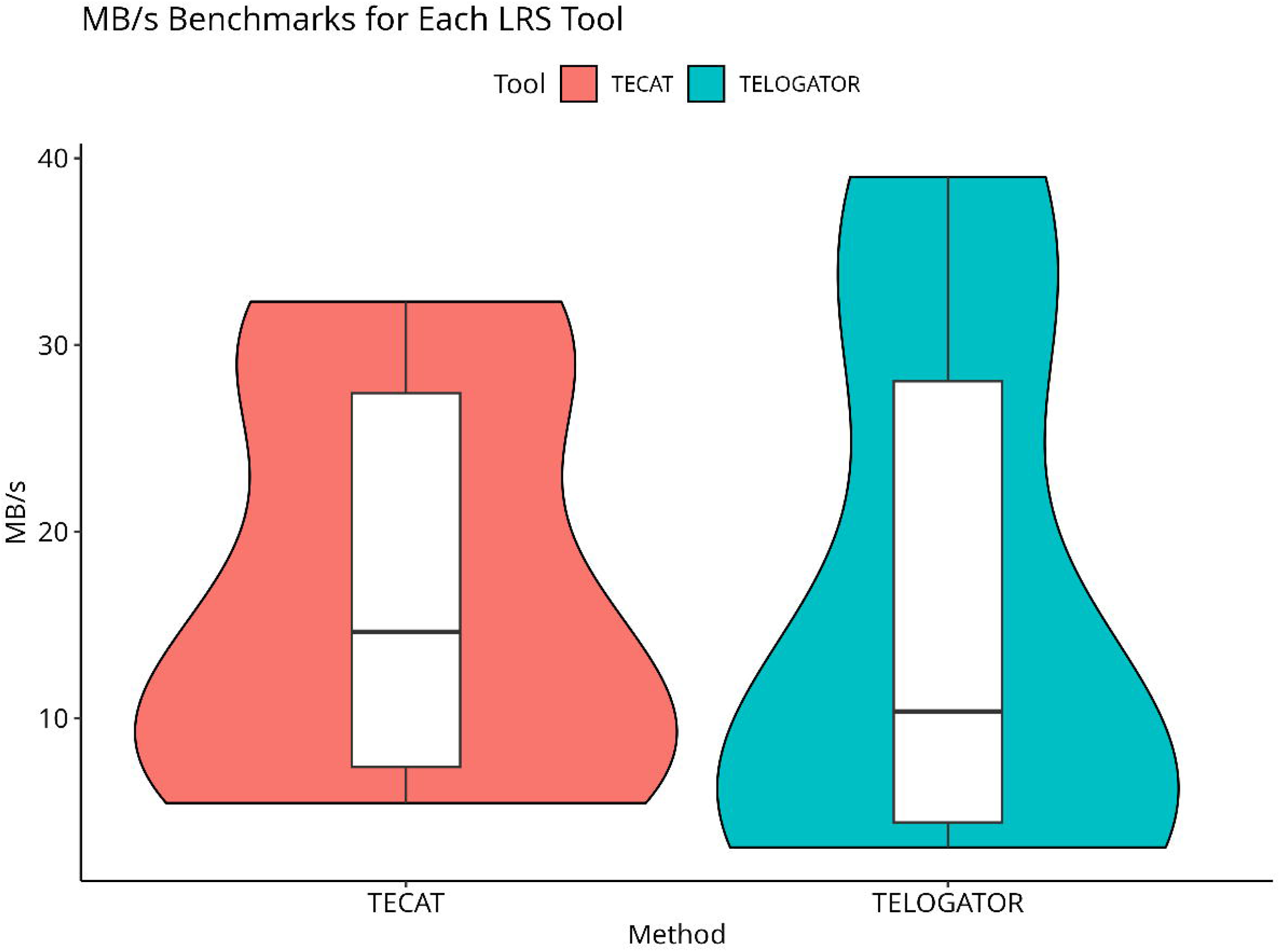
Megabytes processed per second for TECAT and Telogator. The x-axis is the tool that was used while the y-axis is the throughput in megabytes per second.

### 2.3 Chromosome End-Specific Telomere Lengths

Both tools are capable of measuring telomere end-specific lengths. As demonstrated in Fig. 7A and Fig. 7B, these tools perform remarkably well and are able to measure telomere length distributions at the chromosome arm resolution. Upon close inspection of Fig. 7A and Fig. 7B, one is able to see the distribution shape of each chromosome arm closely match each other. This suggests that both tools are capable of measuring relative telomere lengths at the chromosome arm resolution, even if are not absolutely accurate. Both these tools are able to accomplish this feat in a fraction of the time that it would take to accomplish this using traditional methods like TRF or qPCR.

**Fig. 7.**
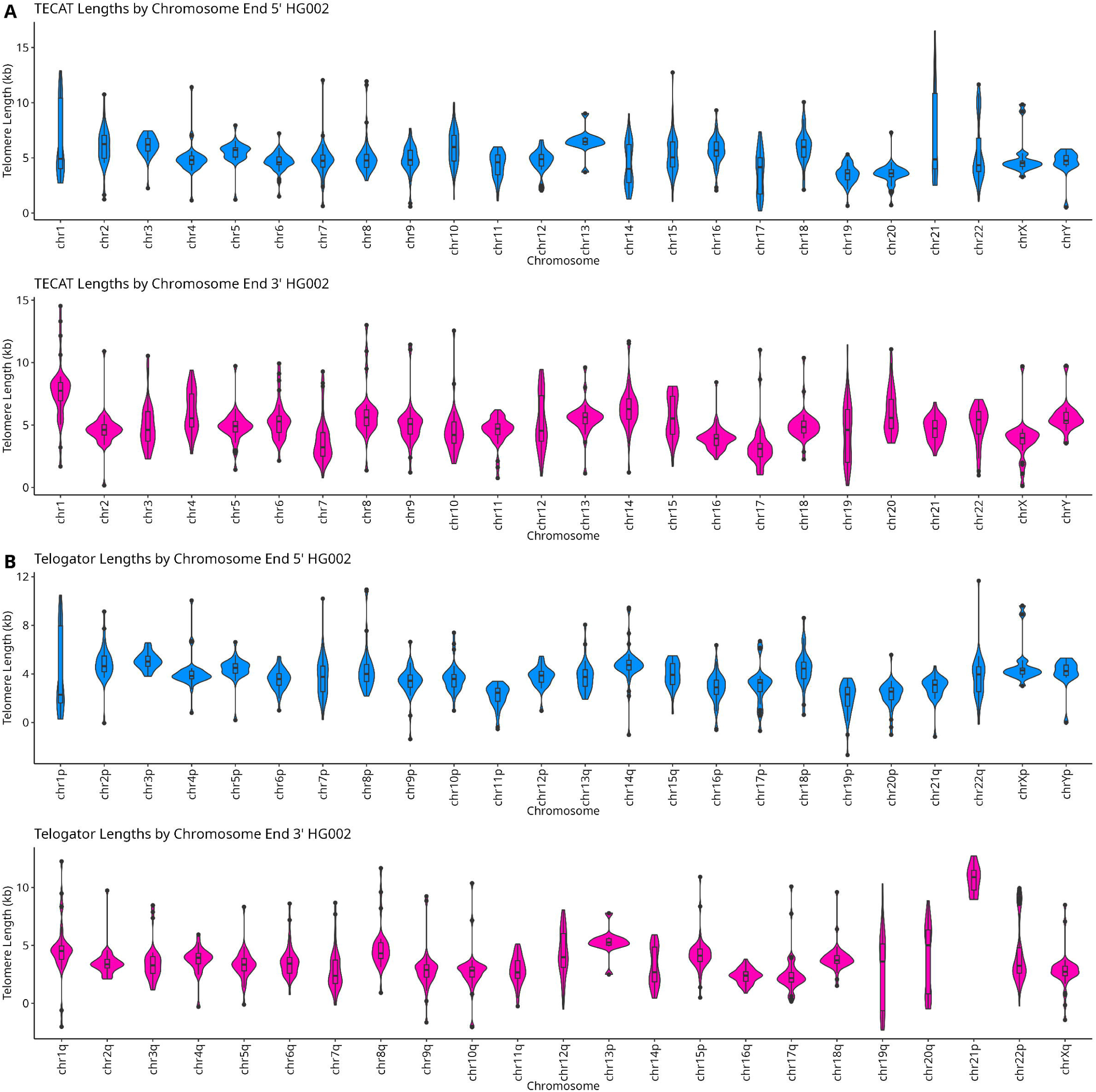
Chromosome end-specific telomere lengths are visualized from TECAT (A) and Telogator (B)

## 3 Methods

### 3.1 Data Collection

The data was all downloaded from the NCBI Sequence Read Archive [36] using the ncbi-toolkit, prefetch and fastq-dump. Table 1 contains all the accession codes necessary to completely reproduce the data set used in this study.

### 3.2 Data Processing

For the short-read data, the data processing was carried out with a custom snakemake [25] pipeline which aligned the reads to the human genome using bwa [21] and then converted to a bam file and sorted the bam file. For the long-read data, all that was required for Telogator and TECAT is a fastq file; therefore, the data was downloaded and dumped into fastq format. TECAT, Telogator [16], and Telseq [24] were then run on the fastq and bam files using a custom bash pipeline to ensure that every sample was run through the same pipeline.

### 3.3 Data Analysis

Data analysis was carried out in R [26] using a custom script which analyzed the resulting data from the processing pipelines. This included plotting length distributions, running Wilcoxon tests and linear regression. The data was then visualized using ggplot2 [29] and cowplot [37].

### 3.4 Data and Code Availability

The code for TECAT can be found at https://github.com/jake-bioinfo/tecat and for Telogator at https://github.com/zstephens/telogator2. The code for the analysis can be found at https://github.com/jake-bioinfo/telo comparative study. The data is available through the Sequence Read Archive (SRA) and the accession codes are available in Table 1.

## 4 Discussion

Measuring chromosome end-specific telomere lengths has been a challenging journey. Now it appears there are at least two tools which are exemplary at measuring chromosome end-specific telomere lengths. The avenues of possible application of this technology are expansive from precision medicine to basic research in mechanisms underlying telomere length dynamics. The chromosome end-specific resolution allows for the generation of all new hypotheses regarding telomere dynamics. The tools are not without their limitations, however, and it is important to note that they are functional, accurate and efficient. Fields in which these tools could be applied to include cancer and ageing research, neruodegenerative diseases research, space exploration, and physical fitness research. Also, these tools could be used in the development of new drugs and therapies. One advantage these tools have over methods where capture is used, is given they can be applied to whole genome long-read sequencing data experimental protocols don’t have to be altered, and these tools could be run on data generated that was not intended for telomere analysis. These tools are at the cutting-edge of long-read sequencing technology and are likely to be the standard for telomere analysis in the future.

## 5 Conclusion

The purpose of this study was to determine what the strengths and weaknesses were for current computational long-read sequencing telomere length analysis tools. The results of this study suggest that Telogator is more sensitive while TECAT is more accurate. TECAT is able to accomplish this in a shorter time span than Telogator as well. This is a great stride forward in telomere length analysis technology, and it is likely that these tools will be the standard for telomere length analysis in the future. The limitations of the study were abundant, but most important, the study was not able to compare any of the results with diseases or aging, due to the 1000 genome project not including any metadata about the patients. Future studies should include this information in order to determine the clinical relevance of these tools. In the future, we plan to use these tools to analyze telomere lengths in a wide variety of patients and conditions which should be able to provide a wealth of information about clinical diseases as well as mechanistic underpinnings of a great deal of diseases.

## Acknowledgements

This work was supported by work performed by Keefer Lin and Pradyun Pulipaka. During their summer research at the Georgia Cancer Center Summer Research Experience for High School Students. They contribute some of the code used in software packages used, wrote some of the documentation for TECAT, and helped with enlightening conversations about the project.

## Declarations

### Ethics approval and consent to participate

Not applicable.

### Consent for publication

Not applicable.

### Materials availability

Not applicable.

### Data availability

SRA accession codes are available in Table 1.

### Author contribution

Jake Reed conceived of the presented idea. Jake Reed developed the theory and performed the computations. Jake Reed and Kevin Coombes verified the analytical methods. Kevin Coombes supervised the findings of this work. Mark Oelkuct developed the shiny app for TECAT. All authors discussed the results and contributed to the final manuscript.

## References

[1] Blackburn, E.H., Epel, E.S., Lin, J.: Human telomere biology: A contributory and interactive factor in aging, disease risks, and protection. Science 350(6265), 1193–1198 (2015) 10.1126/science.aab3389. Publisher: American Association for the Advancement of Science. Accessed 2024-11-20

[2] Lange, T.: Shelterin-Mediated Telomere Protection. Annual Review of Genetics 52, 223–247 (2018) 10.1146/annurev-genet-032918-021921

[3] Shay, J.W., Wright, W.E.: Telomeres and telomerase: three decades of progress. Nature Reviews Genetics 20(5), 299–309 (2019) 10.1038/s41576-019-0099-1. Number: 5 Publisher: Nature Publishing Group. Accessed 2024-11-20

[4] Armanios, M., Blackburn, E.H.: The telomere syndromes. Nature Reviews. Genetics 13(10), 693–704 (2012) 10.1038/nrg3246

[5] Cawthon, R.M.: Telomere measurement by quantitative PCR. Nucleic Acids Research 30(10), 47 (2002). Accessed 2024-11-20

[6] Kimura, M., Stone, R.C., Hunt, S.C., Skurnick, J., Lu, X., Cao, X., Harley, C.B., Aviv, A.: Measurement of telomere length by the Southern blot analysis of terminal restriction fragment lengths. Nature Protocols 5(9), 1596–1607 (2010) 10.1038/nprot.2010.124. Number: 9 Publisher: Nature Publishing Group. Accessed 2024-11-20

[7] Tham, C.-Y., Poon, L., Yan, T., Koh, J.Y.P., Ramlee, M.K., Teoh, V.S.I., Zhang, S., Cai, Y., Hong, Z., Lee, G.S., Liu, J., Song, H.W., Hwang, W.Y.K., Teh, B.T., Tan, P., Xu, L., Koh, A.S., Osato, M., Li, S.: High-throughput telomere length measurement at nucleotide resolution using the PacBio high fidelity sequencing platform. Nature Communications 14(1), 281 (2023) 10.1038/s41467-023-35823-7. Publisher: Nature Publishing Group. Accessed 2024-11-20

[8] Sanchez, S.E., Gu, Y., Wang, Y., Golla, A., Martin, A., Shomali, W., Hockemeyer, D., Savage, S.A., Artandi, S.E.: Digital telomere measurement by long-read sequencing distinguishes healthy aging from disease. Nature Communications 15(1), 5148 (2024) 10.1038/s41467-024-49007-4. Publisher: Nature Publishing Group. Accessed 2024-11-20

[9] Schmidt, T.T., Tyer, C., Rughani, P., Haggblom, C., Jones, J.R., Dai, X., Frazer, K.A., Gage, F.H., Juul, S., Hickey, S., Karlseder, J.: High resolution long-read telomere sequencing reveals dynamic mechanisms in aging and cancer. Nature Communications 15, 5149 (2024) 10.1038/s41467-024-48917-7. Accessed 2024-12-21

[10] Baird, D.M., Rowson, J., Wynford-Thomas, D., Kipling, D.: Extensive allelic variation and ultrashort telomeres in senescent human cells. Nature Genetics 33(2), 203–207 (2003) 10.1038/ng1084. Publisher: Nature Publishing Group. Accessed 2024-11-20

[11] Lai, T.-P., Zhang, N., Noh, J., Mender, I., Tedone, E., Huang, E., Wright, W.E., Danuser, G., Shay, J.W.: A method for measuring the distribution of the shortest telomeres in cells and tissues. Nature Communications 8(1), 1356 (2017) 10.1038/s41467-017-01291-z. Publisher: Nature Publishing Group. Accessed 2024-11-20

[12] Bendix, L., Horn, P.B., Jensen, U.B., Rubelj, I., Kolvraa, S.: The load of short telomeres, estimated by a new method, Universal STELA, correlates with number of senescent cells. Aging Cell 9(3), 383–397 (2010) 10.1111/j.1474-9726.2010.00568.x

[13] Lee, M., Teber, E.T., Holmes, O., Nones, K., Patch, A.-M., Dagg, R.A., Lau, L.M., Lee, J.H., Napier, C.E., Arthur, J.W., Grimmond, S.M., Hayward, N.K., Johansson, P.A., Mann, G.J., Scolyer, R.A., Wilmott, J.S., Reddel, R.R., Pearson, J.V., Waddell, N., Pickett, H.A.: Telomere sequence content can be used to determine ALT activity in tumours. Nucleic Acids Research 46(10), 4903–4918 (2018) 10.1093/nar/gky297. Accessed 2024-11-20

[14] Farmery, J.H.R., Smith, M.L., NIHR BioResource -Rare Diseases, Lynch, A.G.: Telomerecat: A ploidy-agnostic method for estimating telomere length from whole genome sequencing data. Scientific Reports 8(1), 1300 (2018) 10.1038/s41598-017-14403-y

[15] Reed, J., Kirkman, L.A., Kafsack, B.F., Mason, C.E., Deitsch, K.W.: Telomere length dynamics in response to DNA damage in malaria parasites. iScience 24(2), 102082 (2021) 10.1016/j.isci.2021.102082. Accessed 2024-05-31

[16] Stephens, Z., Ferrer, A., Boardman, L., Iyer, R.K., Kocher, J.-P.A.: Telogator: a method for reporting chromosome-specific telomere lengths from long reads. Bioinformatics 38(7), 1788–1793 (2022) 10.1093/bioinformatics/btac005. Accessed 2024-11-18

[17] Montpetit, A.J., Alhareeri, A.A., Montpetit, M., Starkweather, A.R., Elmore, L.W., Filler, K., Mohanraj, L., Burton, C.W., Menzies, V.S., Lyon, D.E., Collins, J.B., Teefey, J.M., Jackson-Cook, C.K.: Telomere Length: A Review of Methods for Measurement. Nursing research 63(4), 289–299 (2014) 10.1097/NNR.0000000000000037. Accessed 2024-11-20

[18] Aubert, G., Hills, M., Lansdorp, P.M.: Telomere length measurement-caveats and a critical assessment of the available technologies and tools. Mutation Research 730(1-2), 59–67 (2012) 10.1016/j.mrfmmm.2011.04.003

[19] Turner, K.J., Vasu, V., Greenall, J., Griffin, D.K.: Telomere length analysis and preterm infant health: the importance of assay design in the search for novel biomarkers. Biomarkers in Medicine 8(4), 485–498 (2014) 10.2217/bmm.14.13

[20] Martin-Ruiz, C.M., Baird, D., Roger, L., Boukamp, P., Krunic, D., Cawthon, R., Dokter, M.M., Harst, P., Bekaert, S., Meyer, T., Roos, G., Svenson, U., Codd, V., Samani, N.J., McGlynn, L., Shiels, P.G., Pooley, K.A., Dunning, A.M., Cooper, R., Wong, A., Kingston, A., Zglinicki, T.: Reproducibility of telomere length assessment: an international collaborative study. International Journal of Epidemiology 44(5), 1673–1683 (2015) 10.1093/ije/dyu191

[21] Li, H., Durbin, R.: Fast and accurate short read alignment with Burrows-Wheeler transform. Bioinformatics (Oxford, England) 25(14), 1754–1760 (2009) 10.1093/bioinformatics/btp324

[22] Li, H., Handsaker, B., Wysoker, A., Fennell, T., Ruan, J., Homer, N., Marth, G., Abecasis, G., Durbin, R., 1000 Genome Project Data Processing Subgroup: The Sequence Alignment/Map format and SAMtools. Bioinformatics 25(16), 2078–2079 (2009) 10.1093/bioinformatics/btp352. Accessed 2025-01-22

[23] Li, H.: Minimap2: pairwise alignment for nucleotide sequences. Bioinformatics (Oxford, England) 34(18), 3094–3100 (2018) 10.1093/bioinformatics/bty191

[24] Ding, Z., Mangino, M., Aviv, A., Spector, T., Durbin, R., UK10K Consortium: Estimating telomere length from whole genome sequence data. Nucleic Acids Research 42(9), 75 (2014) 10.1093/nar/gku181

[25] Molder, F., Jablonski, K.P., Letcher, B., Hall, M.B., Tomkins-Tinch, C.H., Sochat, V., Forster, J., Lee, S., Twardziok, S.O., Kanitz, A., Wilm, A., Holtgrewe, M., Rahmann, S., Nahnsen, S., Koster, J.: Sustainable data analysis with Snakemake. F1000Research (2021). 10.12688/f1000research.29032.1. Accessed 2025-01-22

[26] R Core Team: R: A Language and Environment for Statistical Computing. R Foundation for Statistical Computing, Vienna, Austria (2021). https://www.R-project.org/

[27] Wickham, H.: The Split-Apply-Combine Strategy for Data Analysis. Journal of Statistical Software 40(1), 1–29 (2011)

[28] Xie, Y.: Knitr: A General-Purpose Package for Dynamic Report Generation in R, (2024). https://yihui.org/knitr/

[29] Wickham, H.: Ggplot2: Elegant Graphics for Data Analysis. Springer,(2016). https://ggplot2.tidyverse.org

[30] Kassambara, A.: Ggpubr: ‘ggplot2’ Based Publication Ready Plots, (2023). https://rpkgs.datanovia.com/ggpubr/

[31] Tan, K.-T., Slevin, M.K., Meyerson, M., Li, H.: Identifying and correcting repeat-calling errors in nanopore sequencing of telomeres. Genome Biology 23(1), 180 (2022) 10.1186/s13059-022-02751-6. Accessed 2024-12-19

[32] Tan, K.-T., Slevin, M.K., Leibowitz, M.L., Garrity-Janger, M., Shan, J., Li, H., Meyerson, M.: Neotelomeres and telomere-spanning chromosomal arm fusions in cancer genomes revealed by long-read sequencing. Cell Genomics 4(7) (2024) 10.1016/j.xgen.2024.100588. Publisher: Elsevier. Accessed 2024-12-19

[33] Butler, M.G., Tilburt, J., DeVries, A., Muralidhar, B., Aue, G., Hedges, L., Atkinson, J., Schwartz, H.: Comparison of Chromosome Telomere Integrity in Multiple Tissues from Subjects at Different Ages. Cancer genetics and cytogenetics 105(2), 138–144 (1998) 10.1016/s0165-4608(98)00029-6. Accessed 2025-01-22

[34] Human Genome Assembly GRCh38.p13 -Genome Reference Consortium. https://www.ncbi.nlm.nih.gov/grc/human/data?asm=GRCh38.p13 Accessed 2025-01-22

[35] Shammas, M.A.: Telomeres, lifestyle, cancer, and aging. Current Opinion in Clinical Nutrition & Metabolic Care 14(1), 28 (2011) 10.1097/MCO.0b013e32834121b1. Accessed 2024-12-19

[36] Leinonen, R., Sugawara, H., Shumway, M.: The Sequence Read Archive. Nucleic Acids Research 39(Database issue), 19–21 (2011) 10.1093/nar/gkq1019. Accessed 2025-01-22

[37] Wilke, C.O.: Cowplot: Streamlined Plot Theme and Plot Annotations For ‘ggplot2’, (2024). https://wilkelab.org/cowplot/

